# Cytoskeletal Tubulin Competes with Actin to Increase Deformability of Metastatic Melanoma Cells

**DOI:** 10.1101/2020.02.01.929919

**Authors:** Ghodeejah Higgins, Jade Peres, Tamer Abdalrahman, Muhammad H Zaman, Dirk M Lang, Sharon Prince, Thomas Franz

**Affiliations:** Division of Biomedical Engineering, Department of Human Biology, University of Cape Town, South Africa; Division of Cell Biology, Department of Human Biology, University of Cape Town, South Africa; Department of Biomedical Engineering and Howard Hughes Medical Institute, Boston University, United States of America; Division of Physiological Sciences, Department of Human Biology, University of Cape Town, South Africa; Bioengineering Science Research Group, Engineering Sciences, University of Southampton, UK

**Author notes:** Corresponding author Thomas Franz, Department of Human Biology, Faculty of Health Sciences, University of Cape Town, Private Bag X3, Observatory 7935.

**Keywords:** Cancer, Cell mechanics, Microrheology, Stiffness, Disease stage

## Abstract

The formation of membrane protrusions during migration is reliant upon the cells’ cytoskeletal structure and stiffness. It has been reported that actin disruption blocks protrusions and decreases cell stiffness whereas microtubule disruption blocks protrusion but increases stiffness in several cell types. In melanoma, cell migration is of concern as this cancer spreads unusually rapidly during early tumour development. The aim of this study was to characterise motility, structural properties and stiffness of human melanoma cells at radial growth phase (RGP), vertical growth phase (VGP), and metastatic stage (MET) in two-dimensional *in vitro* environments. Wound assays, western blotting and mitochondrial particle tracking were used to assess cell migration, cytoskeletal content and intracellular fluidity. Our results indicate that cell motility increase with increasing disease stage. Despite their different motility, RGP and VGP cells exhibit similar fluidity, actin and tubulin levels. MET cells, however, display increased fluidity which was associated with increased actin and tubulin content. Our findings demonstrate an interplay between actin and microtubule activity and their role in increasing motility of cells while minimizing cell stiffness at advanced disease stage. In earlier disease stages, cell stiffness may however not serve as an indicator of migratory capabilities.

## 1. INTRODUCTION

Advanced melanoma is a life-threatening cancer that manifests when epidermal melanocytes, pigment producing skin cells, malignantly transform, leading to rapid progression of the disease (Muinonen-Martin *et al*. 2014). When detected early, melanoma can be easily managed and treated (Gladfelter *et al*. 2017). Advanced melanoma is, however, hard to treat and often leads to high mortality rates. During early stage melanoma, cell spread is largely lateral along the epidermis (Gaggioli and Sahai 2007), known as the radial growth phase (RGP). Many tumour genetic features are largely absent during this phase, including metastatic potential and tumour-forming ability, rendering these cancer cells relatively benign. However, as cells transform to the next stage, known as the vertical growth phase (VGP), they acquire pronounced invasion capabilities, where, in addition to lateral spread, they spread vertically, either upwards into the epidermis or downwards invading the basement membrane into the dermis (Gaggioli and Sahai 2007). These cells usually spread in clusters and can form tumours, with only a few cells spreading as individual cells. Finally, VGP cells can transform to the metastatic (MET) stages, where cells characteristically detach from their initial tumour and spread to distant parts of the body where they grow new tumours.

Owing to its characteristic rapid spread during early tumour development, most studies have concentrated on elucidating the molecular and chemical mechanisms that mediate melanoma progression. Despite widespread reports that cells in general alter their mechanical properties to enhance their migratory behaviour (Jonas *et al*. 2011, Pachenari *et al*. 2014), very few studies have focused on the mechanics that enable melanoma cells to spread. Indeed, an inverse correlation between cell stiffness and disease progression have been reported in a variety of cancer types (Gal and Weihs 2012, Omidvar *et al*. 2014, Pachenari *et al*. 2014, Swaminathan *et al*. 2011). In addition, several studies have focused on understanding the cytoskeletal structural properties of cancer cells, such as the role of the actin and tubulin network in facilitating metastatic behaviour. There was general consensus that a reduction in actin filaments (Pachenari *et al*. 2014, Remmerbach *et al*. 2009, Seyedpour *et al*. 2015, Xu *et al*. 2012), but an increase in tubulin (Ferrandina *et al*. 2006), are the molecular mechanisms that enable cells to gain increasing metastatic potential. Pachenari *et al*. (2014) suggest that decreased actin content in high versus low grade colon cancer lines may be the primary molecular basis leading to decreased deformability in metastatic colon cells.

Elucidating the mechanical and structural properties during cancer progression is therefore important and may provide fundamental insights into the biophysical basis underlying uncontrolled proliferation, tumour-forming ability, migration and invasion, and drug resistance, thus leading to the development of more efficient mechanically targeted treatments.

The aim of the current study was to characterise and compare the motility as well as structural and mechanical properties of RGP, VGP and metastatic melanoma cells in 2D *in vitro* conditions.

## 2. MATERIALS AND METHODS

### 2.1. Cell Culture

WM1650 radial growth phase (RGP), ME1402 vertical growth phase (VGP), and WM1158 metastatic (MET) human melanoma cell lines which represent early, intermediate and advanced stages of melanoma, respectively, were cultured and maintained in RPMI 1640 medium (Highveld Biological, Lyndhurst, UK), supplemented with 10% foetal bovine serum (FBS), 100 U/mL penicillin, and 100 μg/mL streptomycin. For WM1650 cells, the medium was supplemented with 200 nM TPA (12-tetradecanoylphorbol 13-acetate) and 200 pM cholera toxin.

Cells were cultured in 25 cm^2^ flasks (Costar, Corning Life Science, Acton, MA) until they reached sub-confluence and at least 48 hrs prior to any experimentation, WM1650 cells were starved of cholera toxin and TPA as these factors promote proliferation and migration respectively.

### 2.2. Cell Morphology Assessment

For morphology assessment, cells were grown until 60% confluence on rigid 35 mm plates (Costar, Corning Life Science, Acton, MA). Brightfield images of single cells were captured and processed using image analysis tools in Image J. At least 21 cells were used for the morphometric analysis. Area (A) and perimeter (P) of the footprint of isolated cells were measured. Cell size and elongation were represented by the area and the circularity (defined as 4πA/P^2^), respectively.

### 2.3. Migration Assays

Migration was measured by conventional two-dimensional (2D) in vitro scratch motility assays. Briefly, RGP, VGP and MET melanoma cells were counted and plated in 24-well plates (Costar, Corning Life Science, Acton, MA) in triplicate. After 24 hrs, confluent monolayers were formed, and a linear wound was then made by scratching through the monolayer using a sterile pipette tip. Cell debris caused by the introduction of the wound was then removed by replacing existing medium with fresh medium. Further, to block the effects of proliferation, Mitomycin C (Sigma, USA) was added to the wells at a final concentration of 2 µg/ml. Mitomycin C acts as a proliferation inhibitor and is often used as an anti-cancer drug (Paz *et al*. 2012). We therefore optimized the final concentration of Mitomycin C to 2 µg/ml since larger concentrations killed the cells. Low concentrations of mitomycin C (less than 4 µg/ml) has been shown to adequately block proliferation while having no effects on cell morphology and migration (Mckenzie *et al*. 2010).

The wound area was imaged at the time of the scratch (0 hr) and thereafter every 2 hrs for 24 hrs. Images were acquired using a phase contrast microscope (Zeiss, Jena, Germany) and the wound area was measured using Image J (Schneider *et al*. 2012). Migration was represented as migrated area (A_0_ – A_*i*_) versus time, where A_0_ and A_*i*_ represent the initial wound area and the wound area at time point *i*, respectively.

### 2.4. Western Blots

Cells were grown in duplicates in 6-well dishes until 60% confluence was reached. Cells located along the diameter of one of the wells were removed using a sterile pipette tip to encourage migration towards the centre, while the second well was left undisturbed. Cells were further incubated for 6 hrs to allow RGP, VGP and MET cells to reach different migratory potentials, and thereafter harvested for western blot analyses.

Briefly, cells were harvested on ice with a protein harvesting buffer containing RIPA (150 mM NaCl, 1% Triton X-100, 0.1% SDS, 10 mM Tris-HCI, pH 7.5, 1% deoxycholate) and 5% protease inhibitor cocktail (Roche, Germany). Whole cell extracts were incubated at 4°C for 30 min on a shaker, centrifuged at 12000 g for 20 min and supernatants collected for subsequent analysis. Total protein concentrations in sample lysates were determined using the bicinchoninic acid (BCA) assay (Pierce, Rockford, IL, USA) in accordance with the manufacturer’s instructions, where bovine serum albumin was used as the standard. To detect target proteins, actin (47 kDa) and β-tubulin (55 kDa), and reference protein, p38 (38 kDa), 20 µg of protein was loaded in 8% SDS-polyacrylamide gels, and proteins were separated at 100 V for 2 hrs. Protein was transferred from onto a Hybond ECL nitrocellulose membrane (Amersham, Biosciences, UK) at 100 V for 1.5 hrs. Membranes were then blocked with 5% milk containing either PBS/0.1% Tween (PBS/T) or TBS/0.1% Tween (TBS/T) for 1 hr at room temperature and probed with appropriate primary antibodies at 4°C overnight. Primary antibodies included monoclonal anti-actin antibody (1:2000) (Abcam, ACTN04-C4), monoclonal anti-β-tubulin antibody (1:500) (Abcam, TU-06) and rabbit polyclonal anti-p38 antibody (Sigma, St. Louis, MO) which was used as a loading control. Membranes were further incubated in peroxidase-conjugated anti-mouse or anti-rabbit secondary antibody (1:5000) (BioRad, Hercules, CA, USA) for 1 hr at room temperature. Finally, proteins were visualized by enhanced chemiluminescence (Pierce, Rockford, IL, USA) using the SuperSignal West Pico Chemiluminescent Substrate Kit (Pierce, Rockford, IL, USA).

The cytoskeletal content was reported as signal intensities of cytoskeletal proteins relative to the signal intensity of the reference protein, and additionally as relative signal intensity normalized to cell size (signal intensity/mm^2^) during natural migratory conditions.

### 2.5. Microrheology

Melanoma cells were grown on 35 mm glass-bottom dishes (Costar, Corning Life Science, Acton, MA) until they reached 60% confluence. To fluorescently label the mitochondria, media was replaced with media containing 200 nM Mitotracker Green Solution (Life Technologies) 30 min prior to microrheology experiments. Cells were then placed in an environmental chamber and maintained at optimal culture conditions of 37°C and 5% CO_2_. During experiments, single cells were located and 120 s time-lapse images (50 ms per frame) of mitochondrial fluctuations were acquired using an inverted confocal microscope with a 63× 1.4 oil immersion objective and a CCD monochrome camera (Zeiss LSM 510, Carl Zeiss, Microimaging, Germany).

Time-lapse images were post-processed into mitochondrial trajectories using TrackMate (Tinevez *et al*. 2017) in the Fiji Image J distribution (Schindelin *et al*. 2012), and the mean-square displacement (MSD) and MSD-dependent power law coefficient α (fluidity) were calculated using custom MATLAB (2017, Mathworks Inc, Natick, MA, USA) algorithms. Cells containing at least 60 mitochondria were considered for analysis, and experiments were repeated once, with an average cell count of eight cells per cell line.

### 2.6. Statistical Analysis

Experiments were repeated at least once on independent days. Data were assessed for normal distribution using the Shapiro-Wilk test. Normally distributed data are presented as mean and standard deviation, *SD*, (in the text) and standard error of mean, *SEM*, (in graphs). Statistical analyses were carried out using one-way ANOVA. Following the detection of statistical significance, post-hoc analysis was done using either Tukey’s HSD test or Games-Howell tests. Data not normally distributed were analysed using Kruskal-Wallis H test followed by Dunn’s test and are presented as median (*Mdn*). Statistical significance was assumed of *p* < .05. All statistical analyses were performed using SPSS for Windows (Version 25, IBM Corp., Armonk, NY).

## 3. RESULTS

### 3.1. Melanoma progression is accompanied by increased cell elongation

Highly invasive cells often undergo molecular and cellular changes to accommodate increased cell dissemination associated with advanced melanoma. We therefore assessed cell morphology as a parameter to facilitate migration in melanoma cells. Here, we hypothesized that advanced melanoma cells require a more elongated cell shape compared to early stage melanoma to enhance cell motility. Figure 1 shows representative images of melanoma cell size and shape across disease progression.

**Figure 1:**
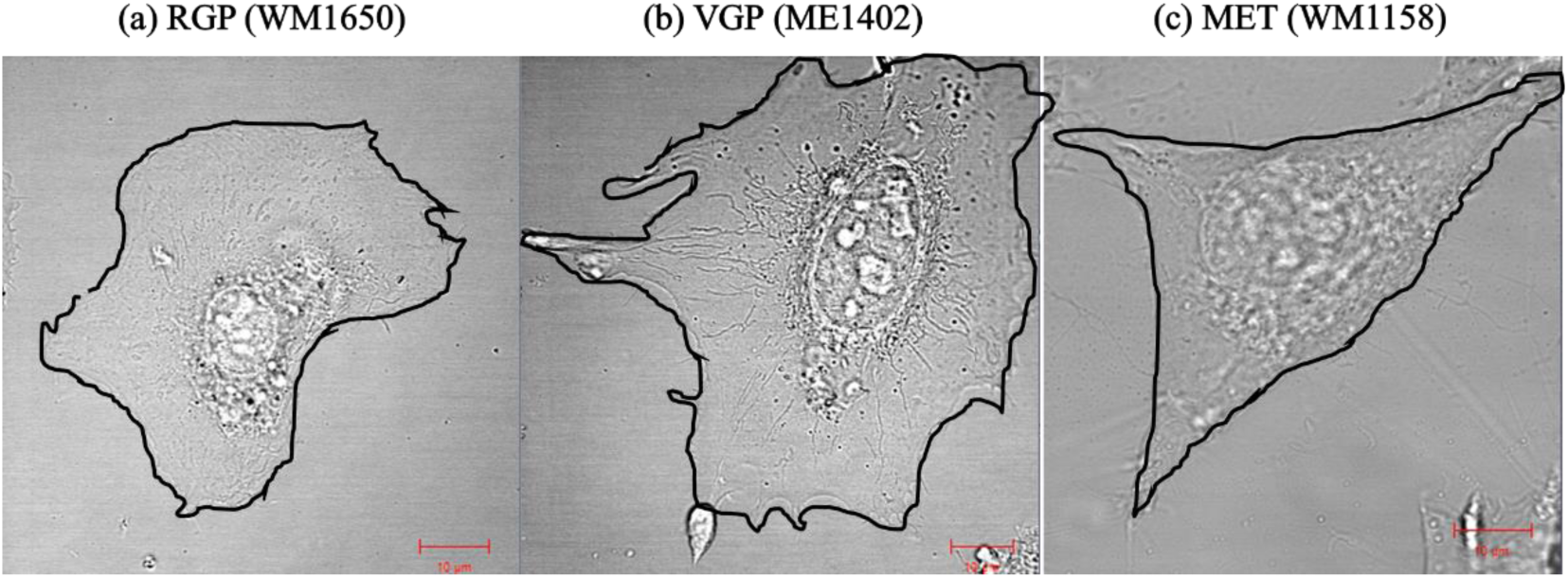
Bright field images showing cell morphology of (a) RGP (WM1650), (b) VGP (ME1402) and (c) MET (WM1158) cells. Scale bars are 10 µm and cell boarders are outlined in black.

When the size (area) of the cells were measured VGP cells (2907 ± 583 µm^2^) were significantly larger than RGP (1513 ± 331 µm^2^, *p* < .001) and MET (1292 ± 315 µm^2^, *p* < .001) cells. The difference in cell size between RGP and MET cells was not significant (*p* = .411). Data are based on n = 2 independent repeats with N = 21 replicates in each group.

The cells were increasingly elongated as the disease progressed. Circularity values indicated that RGP cells (0.61 ± 0.11 µm^2^) were, on average, rounder than both VGP (0.51 ± 0.11 µm^2^, *p* =.004) and MET cells (0.42 ± 0.08 µm^2^, *p* <.001), and VGP cells were rounder than MET cells (*p* = .022). A circularity value of 1 represents a circular shape of a cell and circularity values decreasing towards zero are associated with an increasing elongation of a cell.

### 3.2. Cell motility increases as melanoma disease progresses

To demonstrate that increasingly advanced melanoma cells have higher migration capabilities, we measured cell migration via the scratch motility assay. Figure 2 illustrates typical wound sizes of melanoma cells at 0, 6 and 24 hr time points. Wound sizes of RGP and VGP monolayers were initially (t = 0 hrs) similar (31.96 ± 3.51 and 36.04 ± 8.83 mm^2^, respectively) whereas the same scratch procedure for MET cells resulted in an approximately two-fold wound size (63.14 ± 4.31 mm^2^) (n = 2 independent repeats with N = 3 replicates in each group). Further, RGP cells gradually closed their wounds, decreasing the wound size by 29.1% at the 6 hr time point and only reached half the original wound confluency at the 24 hr timepoint (49.4% reduction of initial wound area). For VGP cells, the initial wound area was reduced by 51.6 and 82.6% after 6 and 24 hr, respectively. MET cells, on the other hand, decreased their initial wound area by 45.4% after 6 hrs and fully closed their wound after 24 hrs. Cell migration was expressed as cell confluence in the wound in which wound closure over time was measured (Figure 2). Cells migrated in accordance with their metastatic potential where they exhibited significant difference in migration upon reaching 6 hrs of wound-induced migration. Indeed, after 6 hrs, RGP cell migration into the wound area (8.87 ± 2.62 mm^2^) was less than both VGP (18.60 ± 6.86 mm^2^, *p* = .037) and MET (28.6 ± 3.33 mm^2^, *p* < .001) cells, and VGP cell migration was less than MET cells (*p* = .033).

**Figure 2:**
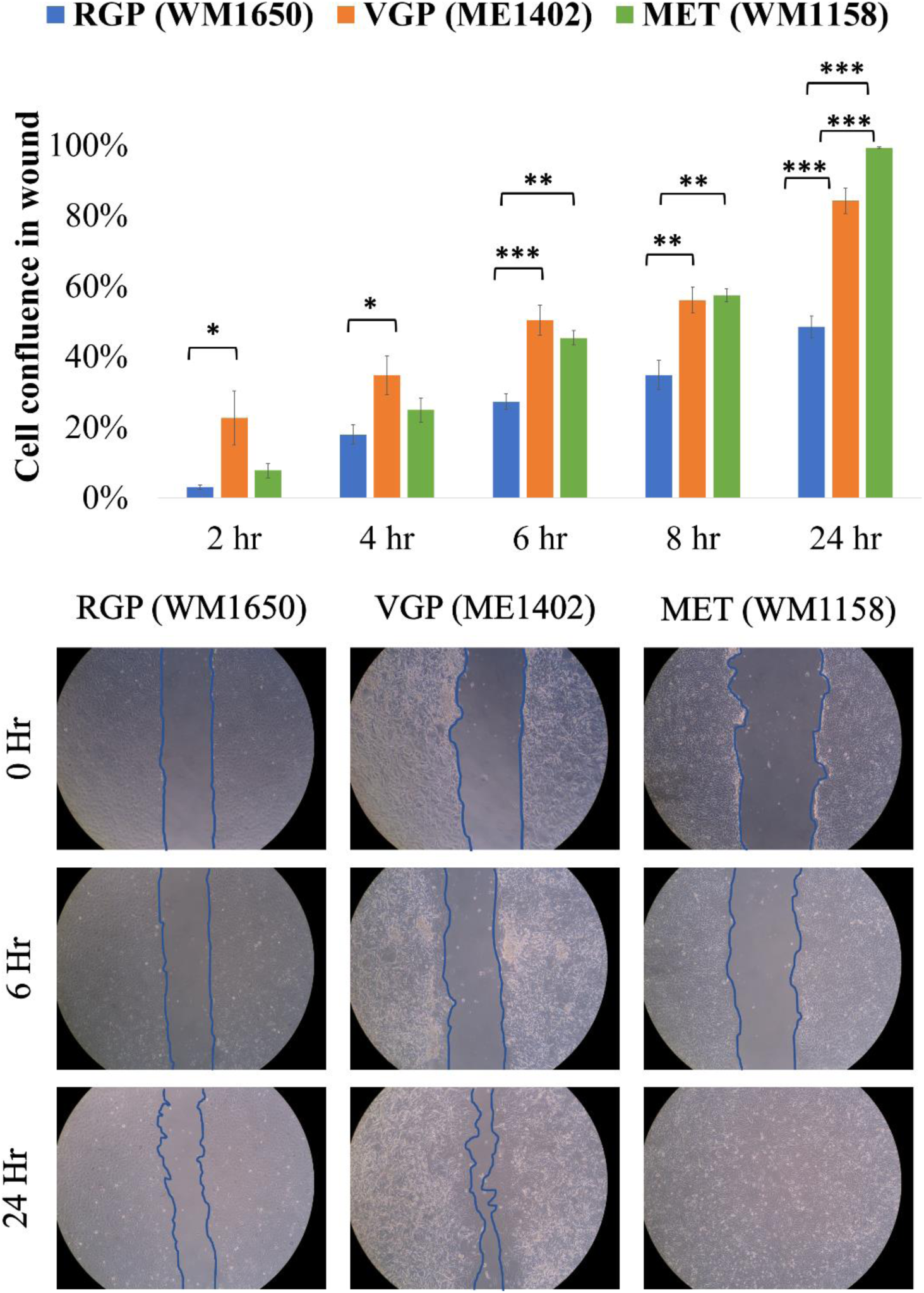
Top: Phase contrast images of melanoma cell monolayers with scratch wounds (10x magnification), at 0, 6 and 24 hrs. Wound edges are outlined by solid lines. Bottom: Graph showing wound closure over time for melanoma cells with increasing metastatic potential. From 6 hrs onwards, RGP cell migration was significantly less than VGP and MET cell migration. Adjusted (* p < .05, ** p < .005, *** p < .0005), Tukey’s HSD test, error bars are SEM for n = 2 independent repeats with N = 3 replicates in each group.

### 3.3. Absolute actin and tubulin contents increase during wound-induced migration

To investigate the structural cytoskeletal content associated with increased cell motility, we compared actin and β-tubulin expression levels in natural and wound-induced migratory conditions in melanoma cells. Figure 3 shows western blot results of actin and β-tubulin expression levels, relative to the reference protein, p38. Irrespective of disease stage, actin and β-tubulin expression were higher for wound-induced migratory conditions compared to natural migration (*p* < .05).

**Figure 3:**
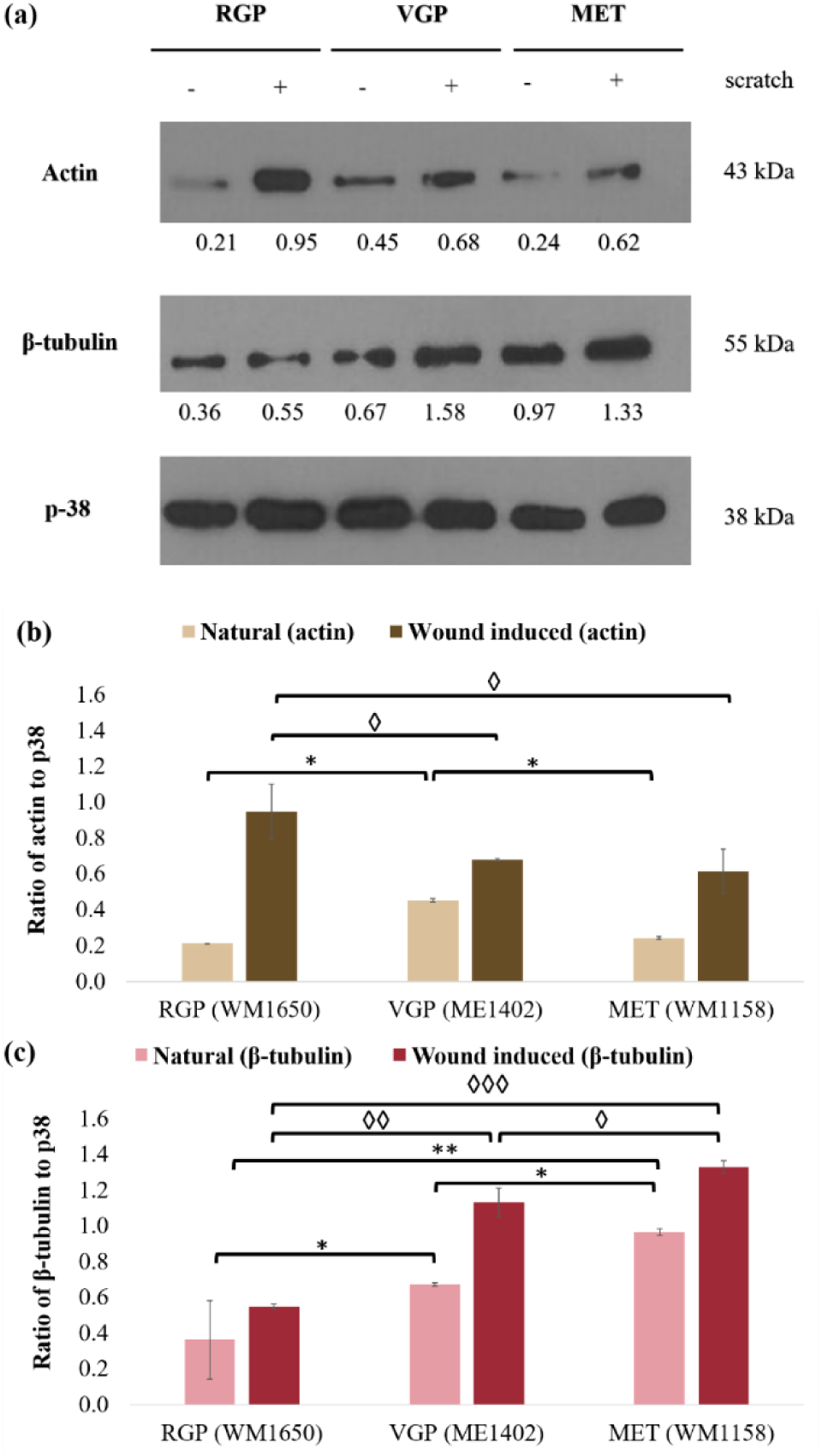
Densimetric readings of western blot experiments. (a) Western blot bands for target proteins (actin and β-tubulin) and loading control (p38). Mean densimetric readings are recorded below each band. The expression of actin and β-tubulin was higher for wound-induced compared to natural migration for all disease stages. Positive (+) and negative (-) symbols indicate with and without a scratch, respectively. Molecular weight of protein is indicated on the right of the bands in kDa. (b) Actin content (normalized to reference protein p38) was higher in VGP cells for natural migration condition whereas it was highest in RGP cells for wound-induced migration. (c) β-tubulin content (normalized to p38) was lowest in RGP cells compared to VGP and MET cells for both natural and wound-induced migration. * ◊ Adjusted p <0.05, Tukey’s HSD Test, error bars are SEM for n = 4 independent repeats with N = 1 replicate.

### 3.4. Levels of tubulin, but not actin, increase with disease progression for natural migration

We hypothesized that with increasing disease progression, melanoma cells will exhibit increased actin and tubulin levels to facilitate the migration process.

Figure 3b shows that during natural migratory conditions, actin levels in VGP cells (0.453 ± 0.012) were higher than in both RGP (0.212 ± 0.004, *p* = .020) and MET (0.243 ± 0.016, *p* = .006) cells. RGP and MET cells, however, had similar actin levels (*p* = .280). During wound-induced migratory conditions, actin levels were higher in RGP cells (1.112 ± 0.322) than in both VGP (0.672 ± 0.016, *p* = .022) and MET (0.603 ± 0.101, *p* = .009) cells. When a wound (“scratch”) was induced, VGP and MET cells had similar actin levels (*p* = .640).

Figure 3c shows that during natural migratory conditions, β-tubulin levels in RGP cells (0.251 ± 0.137) were significantly lower than in VGP (0.672 ± 0.017, *p* = .009) and MET (0.966 ± 0.032, *p* = .001) cells, and β-tubulin levels in VGP cells were significantly lower than in the MET cells (*p* = .041). During wound-induced migratory conditions, a similar trend was observed, where β-tubulin levels in RGP cells (0.550 ± 0.022) were significantly lower than in both VGP cells (1.083 ± 0.070, *p* = .001) and MET cells (1.325 ± 0.044, *p* < .001), and that β-tubulin levels in VGP cells were significantly lower than those in MET cells (*p* = .012).

As shown in Figure 4, when normalized to cell size, actin levels were similar in RGP (140 ± 2.52) and VGP (145 ± 3.98, *p* = .225) cells, and only slightly higher in MET cells (188 ± 8.84) with no statistical differences observed relative to RGP (*p* = .100) and VGP (*p* = .102) cells. The normalised content of β-tubulin was similar in RGP (166 ± 90) and VGP (221 ± 5.72, *p* = .225) cells; however, both RGP and VGP cells had lower normalised β-tubulin content than MET cells (748 ± 25, *p* <.001 for both cases).

**Figure 4:**
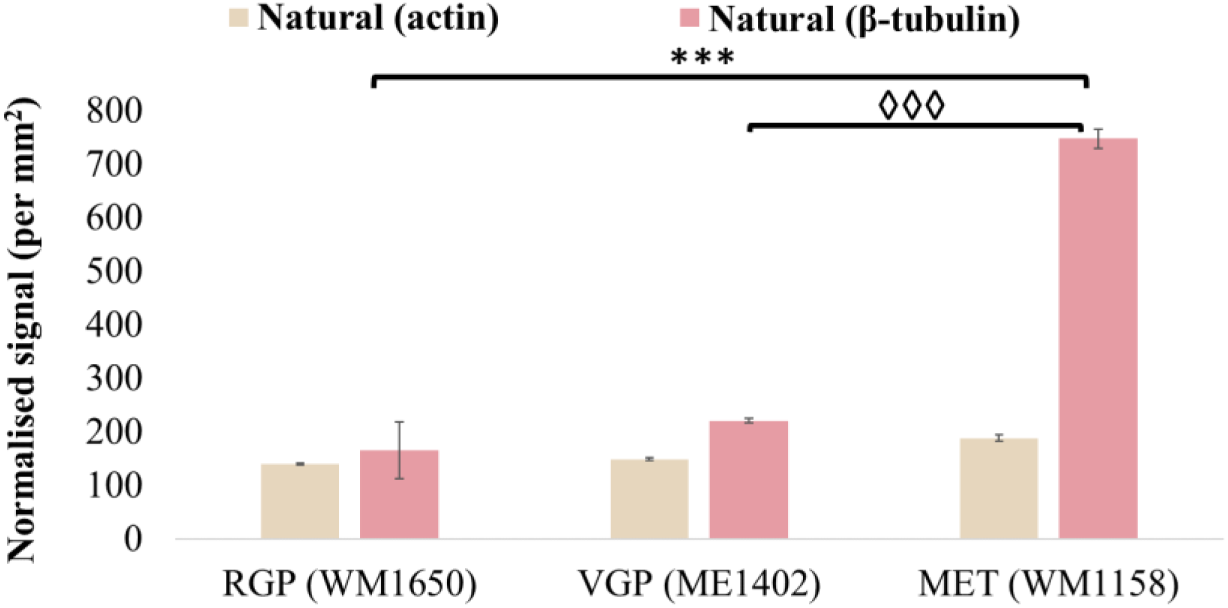
Comparison of cytoskeletal content in natural migration normalized to cell size of melanoma cells. Actin content was similar in all three cell types whereas β-tubulin content was substantially higher in MET cells compared to RGP and VGP cells. * ◊ Adjusted p <0.05, Tukey’s HSD Test, error bars are SEM for n = 4 independent repeats with N = 1 replicate.

### 3.5. Stiffness of RGP and VGP cells is similar and higher than of MET cells

To investigate whether the stiffness of melanoma cells decreases as the disease progresses, we conducted mitochondrial-based particle tracking microrheology. Mitochondrial fluctuations of cells were measured and used to calculate and plot MSD curves as a function of timescales (<Δr^2^ (*t*)>). For viscoelastic materials such as cells and other complex materials, MSD curves follow a power law relationship with <Δr^2^ (*t*)> ∼ t^α^ (Baker *et al*. 2010). The power law exponent α (∂ln <Δr^2^ (*t*)>/∂ln*t*) ranges from α = 0 for elastic (solid-like) materials to α = 1 for viscous (fluid-like) materials with the implication that steeper MSD slopes correspond to increased fluidity and hence decreased stiffness. Figure 5 provides representative images of single cells with stained mitochondria used as tracer particles for microrheology experiments. Generally, RGP cells had spherical mitochondria, VGP cells had filamentous/strand-like mitochondria, and MET cells had mitochondria that possessed a combination of spherical and filamentous structure. Previously, Mak *et al*. (2014) have shown that the MSD of stained mitochondria is comparable to the MSD of ballistically injected nanoparticles with the implication that mitochondria may be used as appropriate tracer particles to measure intracellular mechanics.

**Figure 5:**
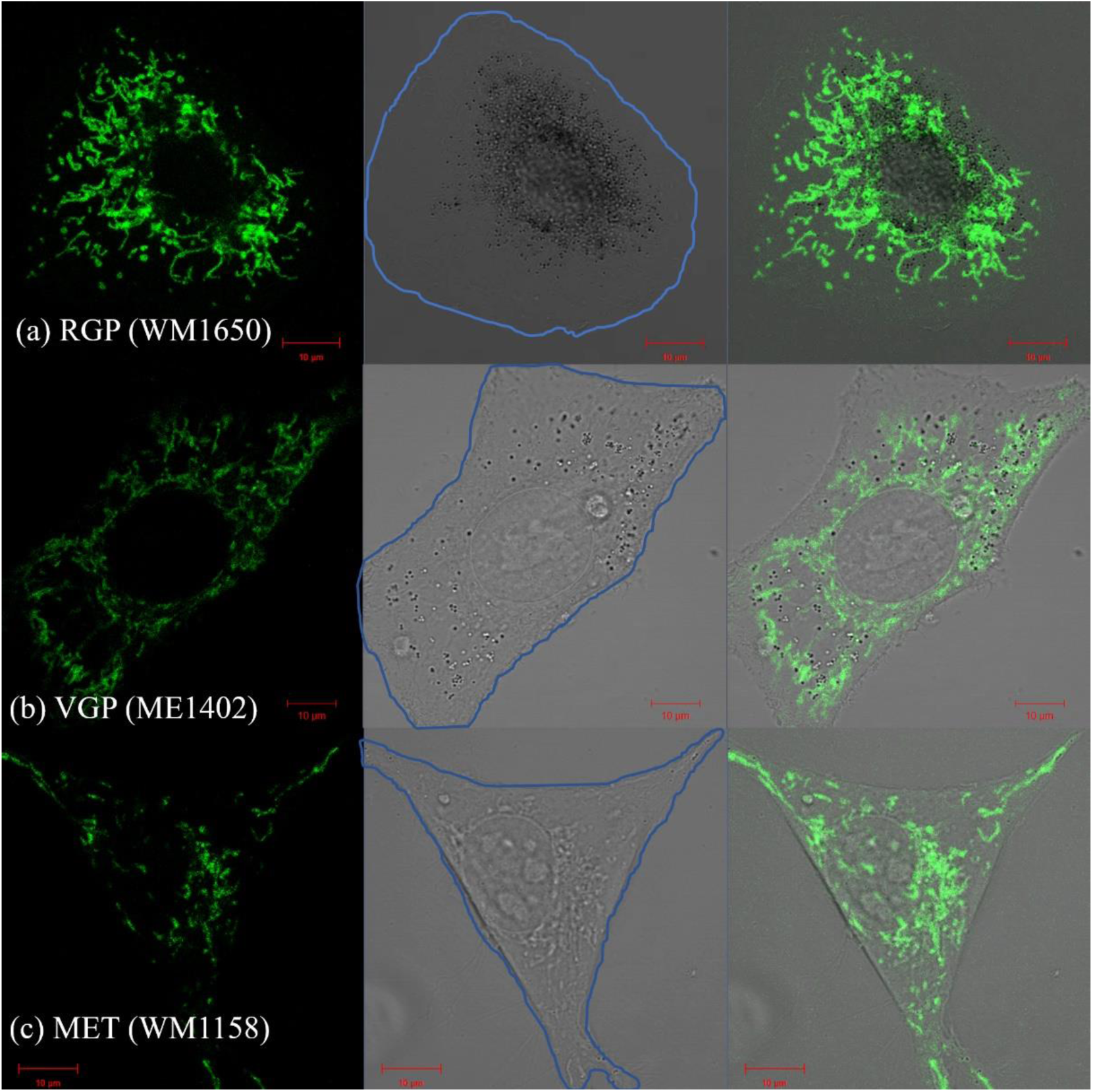
Fluorescent and bright-field images of single cells mitochondrial labelling of (a) RGP, (b) VGP, and (c) MET cells. Mitochondria (green), cell border (blue), scale bars representing 10 µm. Mitochondria in RGP, VGP and MET cells are punctate, filamentous and a combination of punctate and filamentous, respectively.

Figure 6a shows the mean square displacement (MSD) versus delay time for melanoma cells at the different disease stages. The MSD of RGP cells was significantly lower than the MSDs of VGP and MET cells for all delay times. No significant differences in MSD were observed between VGP and MET cells.

**Figure 6:**
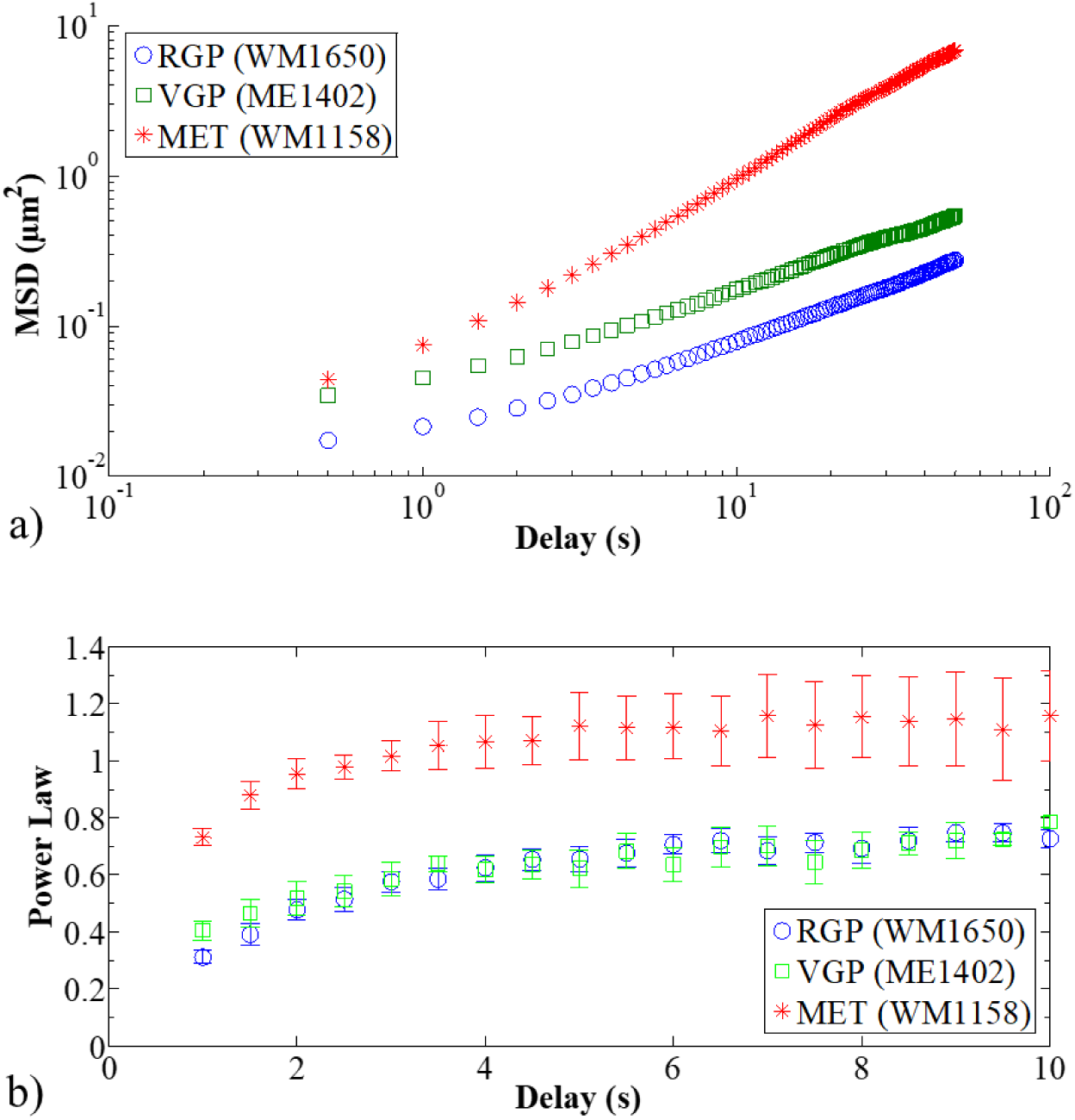
2D mitochondrial MSD and MSD-dependent power law coefficient α (fluidity) versus delay time in melanoma cells. MSD was significantly lower for RGP cells compared to VGP and MET cells (a) whereas the fluidity of MET cells was discernibly higher than that of RGP and VGP cells (b). Error bars indicate SEM for n = 2 independent repeats with N = 8 replicates in each group.

Figure 6b shows the corresponding MSD-dependent power law coefficient α (i.e. fluidity) as a function of delay time. The fluidity of MET cells was discernibly higher than that of RGP and VGP cells for all delay times. Statistically assessing the data at delay times of 1, 2, 5 and 10 s, significant differences between MET and both RGP and VGP were only found at 5 s (*p* = .043 and *p* = .021, respectively). For 1, 2 and 10 s, differences were statistically significant between MET and RGP (*p* = .003, *p* = .012 and *p* = .016, respectively) but not VGP (*p* = .412, *p* = .094 and *p* = .475, respectively).

## 4. DISCUSSION

In this study, we characterised the motility, morphology and mechanics of human melanoma cells at three different disease stages in order to investigate whether a change in mechanical and structural properties enable the cells to increase their metastatic capabilities.

MET cells exhibited a 44.6% higher intracellular fluidity (i.e. deformability) than RGP and VGP cells, whereas the fluidity of VGP cells was similar (8.3% higher) to that of RGP cells. These results agree with many studies (Fraldi *et al*. 2015, Gal and Weihs 2012, Guck *et al*. 2005), which indicate that more motile cells in an advanced disease stage are more deformable than less motile cancer cells at earlier disease stages.

Our finding of similar intracellular fluidity for RGP and VGP cells based on microrheology, however, does not align with previous reports of significant decrease in stiffness of melanoma cells from RGP to VGP stage measured with atomic force microscopy (Weder *et al*. 2014). It should be noted, though, that the authors concluded that atomic force microscopy may not be suitable for staging melanoma.

Whereas changes in motility have been associated with changes in cellular stiffness in melanoma cells (Watanabe *et al*. 2012), we observed a significantly higher absolute migrated area of VGP cells compared to RGP cells (Figure 2) despite the similar intracellular fluidity at the two disease stages. However, normalizing the migrated area to cell size leads to similar motility for VGP and RGP cells. As such, the increased motility of VGP cells in our study is a result of the increased cell size rather than decreased cell stiffness.

In natural migratory conditions, actin levels increased from RGP to VGP cells by 53.3% and decreased by 46.4% from VGP to MET cells. Considering that increased actin leads to increased cellular stiffness (Grady *et al*. 2016, Mak *et al*. 2014, Pachenari *et al*. 2014), the melanoma cells would be expected (a) to stiffen during progression from RGP to VGP disease stage which we, however, did not observe.

Our results indicated that actin increase may be related to cell size. VGP cells were approximately twice the size of RGP and MET cells. Likewise, we observed that VGP cells exhibited a two-fold absolute content of actin compared to RGP and MET cells.

However, when normalizing the actin content to cell size, actin levels were similar for all three disease stages. This suggests that despite increasing motility with increasing disease stage, the cells generate only the amount of cytoskeletal actin needed, but not more, for their structural support. More importantly, normalized cell-substrate adhesion area and traction forces required to displace the cell in the natural migratory state may be similar.

Our results suggest that the reduction of actin content from VGP to MET cells is not a direct consequence of the increase in metastatic properties with disease progression. Rather, the decreased actin level is linked to the smaller cell size that reduces the structural support required by the cell and which is provided by actin.

During wound-induced migration, actin levels decreased from RGP to VGP disease stage by 65% and were similar for VGP and MET cells, with only ≈10% decrease. During wound induced migration, cells need to drastically migrate to close the wounds. Only very few studies have described the mechanics involved in cell migration during melanoma progression (Watanabe *et al*. 2012, Weder *et al*. 2014). Limited data on the change in cytoskeletal protein concentration and cytoskeletal dynamics during disease progression are reported for colon and ovarian cancer (Pachenari *et al*. 2014, Xu *et al*. 2012) but not for melanoma. In line with Pachenari *et al*. (2014), we observed a marked decrease in actin as melanoma cells progressed from relatively benign RGP to more invasive VGP and MET stages for wound-induced migration. However, we did not observe the same for natural migratory conditions. We suggest that in advanced melanoma cells in their natural migratory state, actin levels are less dependent on cell size and migratory capabilities, regardless of the difference in metastatic or invasive potential between VGP and MET.

Tubulin content was found to be proportional to disease progression for both natural and wound-induced migratory conditions. This indicates that regardless of the migratory condition (natural or wound-induced), cells require structural fortification via microtubules to enhance their migratory behaviour. It is noteworthy to mention that although no statistical differences in normalized actin content were observed between the cell lines, MET cells produced slightly more actin compared to RGP and VGP cells. We suggest that the increased action production is rooted in the need of MET cells to be highly motile. However, an increase in the actin content contributes to an increased cell stiffness and thereby diminishes the cell’s capabilities to pass through capillaries with luminal diameters much smaller than the dimension of the cell. Microtubules may indirectly affect cell stiffness by attenuating or counteracting the stiffening effect of actin filaments.

The present method of reporting structural proteins disregards spatial organization of actin which is equally important in determining cell stiffness. Future work therefore should incorporate experimental techniques that provide data of amount of actin as well as its organization. Furthermore, the present study provides the total actin and tubulin concentration after considerable migration has occurred. More sophisticated techniques need to be employed to distinguish between, or decouple, the contributions of actin and tubulin to motility during each phase of the migration process.

This study highlighted mechanical properties indicative of different migration properties of melanoma cells in 2D environments. Further research is required to demonstrate the effects of cancer cell mechanics in physiologically more realistic 3D environments.

It is important to note that cells were used at a 60 % confluence for all experiments, except for the migration assay. However, the aim of measuring collective cell migration via the 2D scratch assay was not to quantitatively compare migration against morphology or cell stiffness. Rather, migration experiments served as preliminary work to determine the timepoint at which cells migrated different to each other for western blot experiments, i.e. 6 hrs after wound-induced migration. Future research could include single cell migration experiments and correlation with single cell morphology and stiffness to extend the findings of this study.

## 5. CONCLUSION

Advanced melanoma manifests as a result of the malignant transformation of melanocytes which further undergoes a rapid stepwise progression from benign radial growth phase to advanced vertical growth phase and metastatic stages. Despite several treatment strategies, such as chemotherapy, surgery and radiation therapy, survival rates of humans diagnosed with advanced melanoma remain poor (Sulaimon and Kitchell 2003). To develop more efficient targeted therapies, it is therefore vital to understand the collective molecular and mechanical mechanisms that mediate cancer metastasis. However, mechanical properties of melanoma cancer cells have been largely understudied, particularly during disease progression. We therefore investigated and demonstrated the impact of intracellular mechanics, such as cell stiffness and cytoskeletal content, associated with different stages in melanoma progression. These results are valuable and open the doors to further studies to develop new cancer therapies that not only targets molecular pathways, but conjointly targets mechanical properties that may lead to invasive cancer behaviour.

## ACKNOWLEDGEMENTS

We acknowledge the technical assistance of Mrs. Susan Cooper of the Confocal and Light Microscope Imaging Facility of the University of Cape Town. The confocal microscope used in this work was purchased with funds from the Wellcome Trust (grant reference number 108473/Z/15/Z and the National Research Foundation (NRF) (grant reference number UID 93197).

## DATA

Data presented in this article are available on ZivaHub under http://doi.org/10.25375/uct.11764050.

